# Identification of ACE2 modifiers by CRISPR screening

**DOI:** 10.1101/2021.06.10.447768

**Authors:** Emily J. Sherman, Carmen Mirabelli, Vi T. Tang, Taslima G. Khan, Andrew A. Kennedy, Sarah E. Graham, Cristen J. Willer, Andrew W. Tai, Jonathan Z. Sexton, Christiane E. Wobus, Brian T. Emmer

## Abstract

SARS-CoV-2 infection is initiated by binding of the viral spike protein to its receptor, ACE2, on the surface of host cells. ACE2 expression is heterogeneous both *in vivo* and in immortalized cell lines, but the molecular pathways that govern ACE2 expression remain unclear. We now report high-throughput CRISPR screens for functional modifiers of ACE2 surface abundance. We identified 35 genes whose disruption was associated with a change in the surface abundance of ACE2 in HuH7 cells. Enriched among these ACE2 regulators were established transcription factors, epigenetic regulators, and functional networks. We further characterized individual cell lines with disruption of *SMAD4, EP300, PIAS1*, or *BAMBI* and found these genes to regulate ACE2 at the mRNA level and to influence cellular susceptibility to SARS-CoV-2 infection. Collectively, our findings clarify the host factors involved in SARS-CoV-2 entry and suggest potential targets for therapeutic development.

## INTRODUCTION

ACE2 plays a critical role in SARS-CoV-2 infection by serving as the cellular receptor for viral entry (Zamorano Cuervo and Grandvaux, 2020). Inhibition of endogenous ACE2 disrupts SARS-CoV-2 entry into permissive cell lines while heterologous expression of ACE2 in non-permissive cell lines renders them susceptible to infection (Hoffmann et al., 2020; Ou et al., 2020; Walls et al., 2020). Transgenic expression of human ACE2 sensitizes mice to SARS-CoV-2 infection with recapitulation of pathologic hallmarks of COVID-19 (Bao et al., 2020; Jiang et al., 2020; Winkler et al., 2020).

Given its critical role in SARS-CoV-2 infection, the interaction between the viral spike protein and ACE2 is an attractive target for therapeutic development. Vaccines against the spike protein are broadly efficacious in reducing the number and severity of COVID-19 infections (Klasse et al., 2021), but the rapid evolution of SARS-CoV-2 raises concern for the potential of SARS-CoV-2 variants to escape immunity induced by either vaccines or prior infection (Cele et al., 2021; Garcia-Beltran et al., 2021; McCarthy et al., 2021). As an alternative strategy, disruption of host ACE2 may similarly prevent SARS-CoV-2 infection in a manner that is less susceptible to viral evolution. ACE2-targeted therapies may have broader clinical applications, as ACE2 also serves as the cellular receptor for other respiratory viruses such as SARS-CoV-1 (Li et al., 2003) and HCoV-NL63 (Hofmann et al., 2005). Additionally, ACE2 is an important physiologic regulator of the renin-angiotensin and kallikrein-kinin systems, and its dysregulation has been implicated in pulmonary and systemic hypertension, cardiac fibrosis, atherosclerosis, and acute respiratory distress syndrome (Gheblawi et al., 2020). However, no ACE2-targeted therapies have been clinically approved and their development is limited by uncertainty in the molecular pathways that regulate ACE2.

Since the onset of the COVID-19 global pandemic, several studies have applied single-cell RNA-seq to examine *ACE2* expression in tissues of humans and animal models (Menon et al., 2020; Sungnak et al., 2020; Ziegler et al., 2020; Zou et al., 2020). These studies have consistently found *ACE2* mRNA expression to be heterogeneous among cell types within a given tissue. *ACE2*-expressing cells include alveolar type 2 cells in the lung, goblet cells in the nasopharynx, absorptive enterocytes in the gut, and proximal tubular epithelial cells in the kidney. Even within a given cellular subtype, *ACE2* expression is heterogeneous, with only ∼1-5% of lung AT2 cells, for example, containing detectable A*CE2* mRNA. Prior investigations have identified two distinct promoters for full-length *ACE2* which vary across tissues in their relative usage (Pedersen et al., 2013), as well as a cryptic promoter driving expression of an interferon-responsive truncated *ACE2* isoform (Ng et al., 2020; Onabajo et al., 2020). Other studies have identified putative transcription factor binding sites and epigenome signatures associated with the *ACE2* locus (Beacon et al., 2020; Chlamydas et al., 2020) as well as transcriptome profiles associated with *ACE2* expression (Barker and Parkkila, 2020; Feng et al., 2020).

Importantly, single cell RNA-seq approaches have intrinsic limitations for low abundance transcripts (Saliba et al., 2014), and most studies of *ACE2* mRNA have lacked validation at the protein level. Investigations of ACE2 protein expression have been relatively limited and confounded by uncertain specificity of different commercial ACE2 antibodies. Recently, we engineered *ACE2*-overexpressing and *ACE2*-deleted cell lines and performed systematic testing of a panel of commercial antibodies by flow cytometry, finding only 2 of 13 to exhibit specificity and sensitivity for ACE2 surface protein (Sherman and Emmer, 2021). Unexpectedly, we found that multiple isogenic cell lines demonstrated heterogeneity of endogenous ACE2 expression, suggesting that they may serve as a simplified model to dissect the molecular pathways that govern ACE2 expression. To this end, we now report the findings of our high-throughput CRISPR screens for modifiers of endogenous ACE2 surface abundance in HuH7 cells. We identified 35 previously unrecognized ACE2 regulators, which we then analyzed for molecular functions, genetic interactions, and influence on viral infection. Arrayed single gene validation studies confirmed the ACE2 regulatory effect for 18 of 20 genes tested, enabled a more detailed characterization of a subset of selected genes, and demonstrated the relevance of these ACE2 modifiers to SARS-CoV-2 infection.

## RESULTS

### CRISPR screen for ACE2 modifiers

To identify functional regulators of ACE2 surface abundance, we first defined a list of candidate genes from several different sources (Figure 1A, Supplemental Table 1). These included modifiers of SARS-CoV-2 cytopathic effect in recently reported CRISPR screens (Daniloski et al., 2021; Gordon et al., 2020; Heaton et al., 2020; Hoffmann et al., 2021; Wei et al., 2021), genes in proximity to human GWAS loci associated with COVID-19 susceptibility (Ganna, 2021) (Supplemental Table 2), genes we previously identified by RNA-seq whose correlation was associated with *ACE2* expression in sorted HuH7 cells (Sherman and Emmer, 2021), candidates identified in our own pilot genome-wide CRISPR screens for ACE2 abundance (Figure S1, Supplemental Tables 3-4), and a set of hypothesis-driven manually selected genes. In total, we targeted 833 genes with 15 gRNA per gene. Amplicons of the pooled gRNA sequences were inserted into the pLentiCRISPRv2 construct (Sanjana et al., 2014) and the diversity and representation of the resulting plasmid pool were verified by deep sequencing.

**Figure 1.**
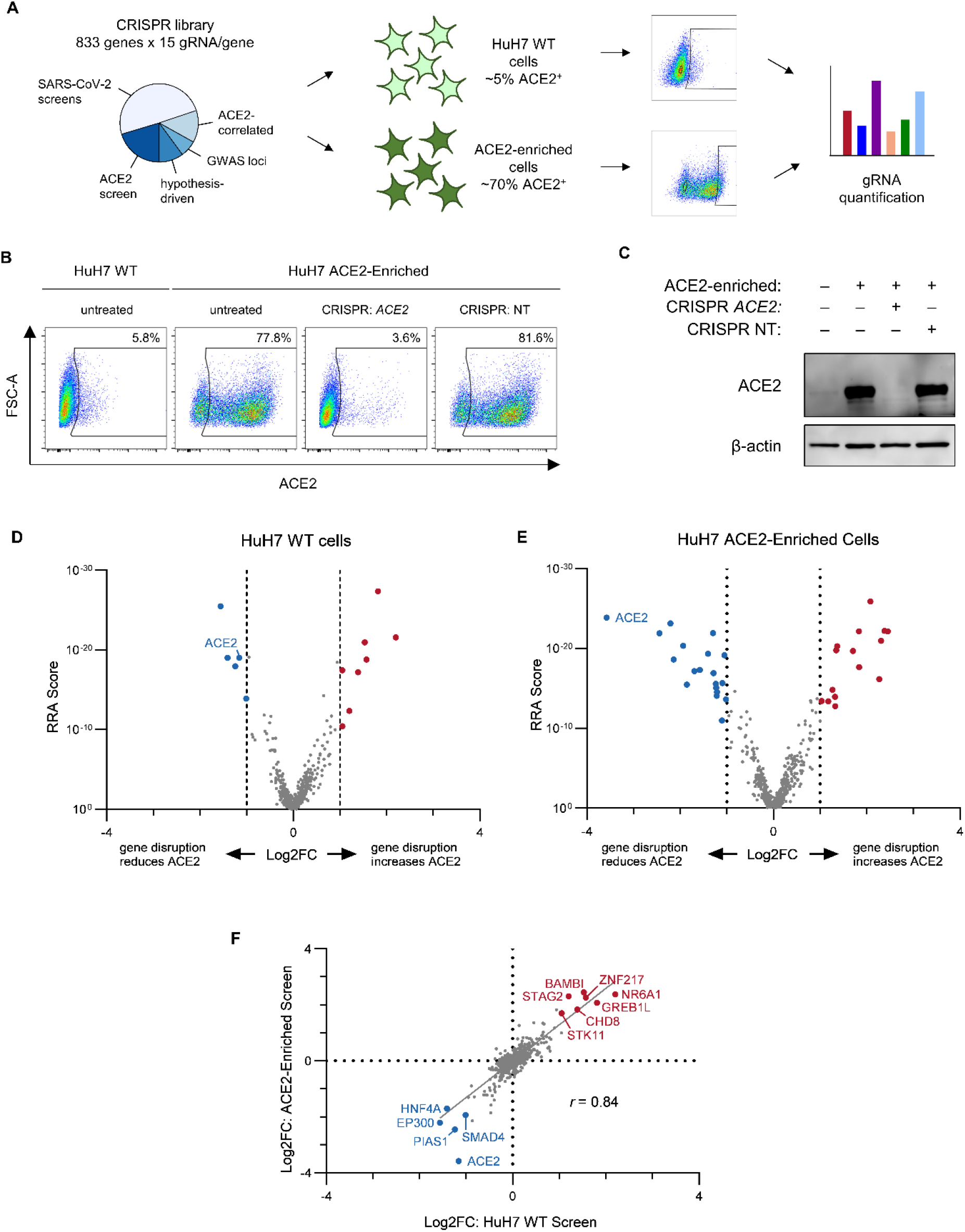
CRISPR screen for ACE2 modifiers. (A) Schematic of ACE2 CRISPR screening strategy, with design and synthesis of a high-resolution focused CRISPR library of candidate ACE2 modifiers used to mutagenize in parallel wild-type or ACE2-enriched HuH7 cells, which were then selected by FACS based on ACE2 surface abundance and gRNA representation in each library quantified by massively parallel sequencing. (B) Flow cytometry plots of ACE2 surface abundance for wild-type or serially ACE2-enriched HuH7 cells, with or without transduction of a lentiviral CRISPR construct with a *ACE2*-targeting or control nontargeting (NT) gRNA. ACE2-positive gates were established on unstained wild-type cells. (C) Immunoblot for ACE2 and β-actin of lysates prepared from the same cell populations as in (B). (D-E) Volcano plots of MAGeCK Robust Rank Aggregation scores relative to gene-level gRNA log2 fold-change for each gene tested in the secondary library in screens of ACE2 surface abundance in HuH7 wild-type (D) or ACE2-enriched (E) cells. Genes with FDR <0.05 and absolute log2 fold-change >1 are highlighted, with positive regulators in blue and negative regulators in red. (F) Correlation of gene-level aggregate gRNA log2 fold-change between the independent secondary screens of HuH7 wild-type and ACE2-enriched cells. Genes identified in both screens with FDR <0.05 and absolute log2 fold-change >1 are highlighted and annotated.

We independently screened both HuH7 wild-type cells, in which ∼3-5% of cells express detectable ACE2, and HuH7 cells derived by serial enrichment with 3 rounds of FACS to contain ∼60-70% ACE2-positive cells (Figures 1B, C). Cells were then transduced at >200X coverage with the customized CRISPR library, passaged for 14 days to allow for target gene mutagenesis and turnover of residual protein, and sorted by flow cytometry into selected populations. In the screen of wild-type cells, the ∼3-5% of ACE2-positive cells were collected in one population and the median ∼10% of ACE2-negative cells were collected in another. In the screen of ACE2-enriched cells, the top ∼10% of cells with the greatest ACE2 abundance were collected along with the median ∼10% of ACE2-negative cells. The relative abundance of every gRNA in each population was quantified by deep sequencing. In accordance with the depth of library coverage in this screen, we found >99.9% library representation in each sorted cell population with minimal skewing of gRNA representation (Figure S3A, B). As expected, *ACE2* itself was identified among the top positive regulators of ACE2 abundance in both screens (Figure S3C, D), while gRNAs targeting many other genes without a known role in ACE2 regulation exhibited significant enrichment or depletion in sorted populations (Figure 1D, E, Supplemental Tables 5, 6). In total, we identified 19 high-confidence positive regulators and 16 high-confidence negative regulators of ACE2 surface abundance in HuH7 cells (FDR<0.05, absolute value of log2 fold-change >1). Supporting the reproducibility of the screen results, there was a very high degree of concordance between the results of the independent screens of wild-type and ACE2-enriched cells (Figure 1F, *r* = 0.84).

### Arrayed validation of ACE2 modifiers

To validate our CRISPR screen results, we generated single gene CRISPR-targeting lentivirus constructs for 21 of the top-scoring hits from either screen and tested whether single gene disruption affected surface ACE2 levels. We found one gene, *UROD*, to be a false positive of the screen due to its disruption causing increased cellular autofluorescence in the detection channel of the Alexa Fluor 647-conjugated secondary antibody. *UROD*-targeted cells exhibited increased fluorescence in this channel even when the conjugated secondary antibody was omitted (Figure S5A), and replacement of this secondary antibody with an Alexa Fluor 488-conjugated secondary antibody abrogated the difference in ACE2 signal (Figure S5B). These findings are consistent with the known role of UROD in decarboxylating uroporphyrinogen-III, as cells with accumulation of uroporphyrin have been shown to exhibit increased fluorescence between 620-680 nm (Schneckenburger et al., 1989). Of the remaining genes, we confirmed a significant effect of single gene CRISPR-targeting on ACE2 surface staining for 18 of 20 genes in either HuH7 wild-type or ACE2-enriched cells (Figures 2, S6, S7; Supplemental Table 7), with none of these genes showing increased autofluorescence and each replicating the change in ACE2 abundance with an Alexa Fluor 488-conjugated secondary antibody. Consistent with the stronger enrichment and depletion of hits in the CRISPR screen of ACE2-enriched cells relative to wild-type cells, validation testing of these ACE2-enriched cells also exhibited increased power to detect significant changes in ACE2 surface abundance (Supplemental Table 7).

**Figure 2.**
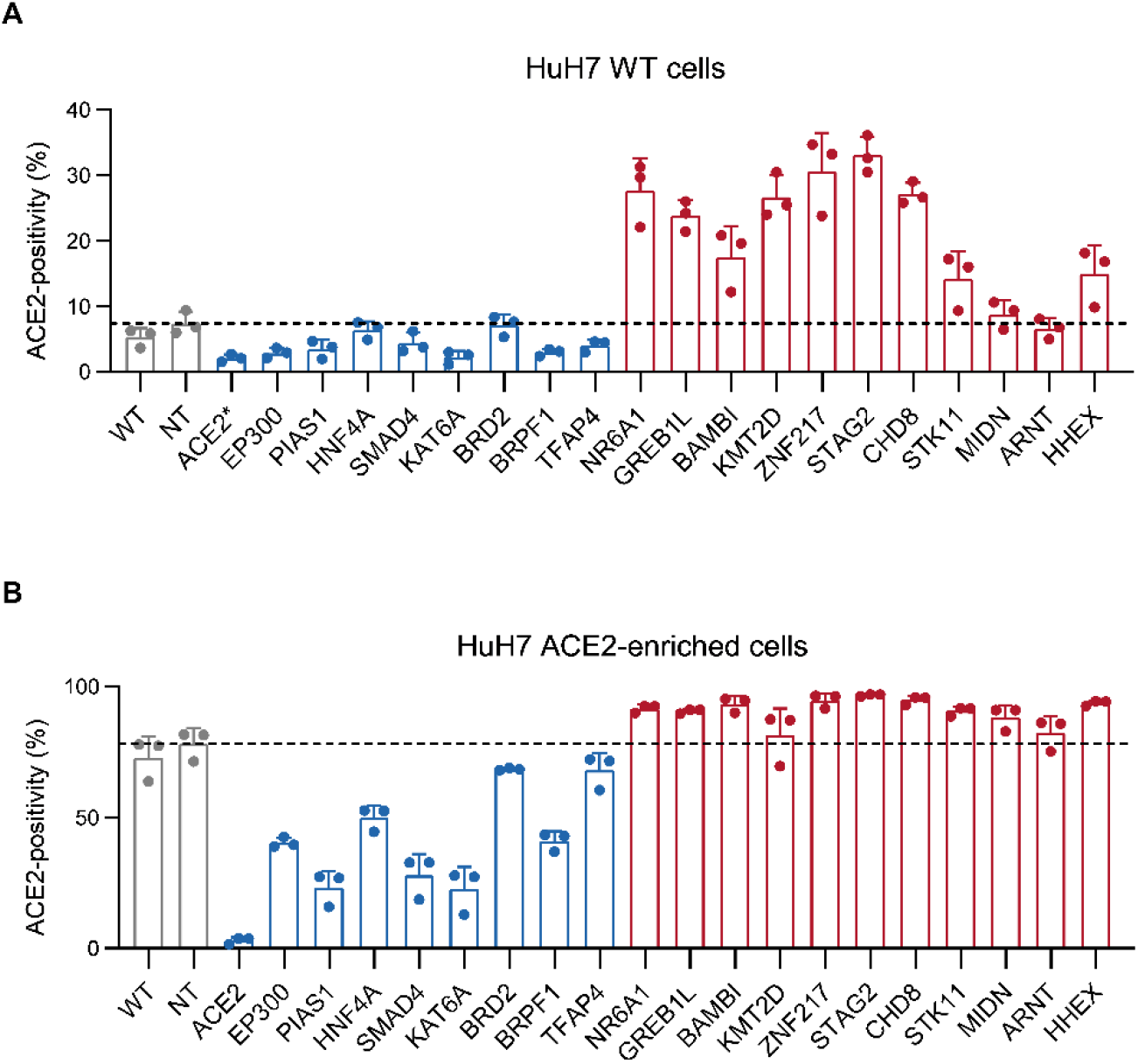
Arrayed validation of ACE2 modifiers. (A-B) Percent ACE2-positive cells for HuH7 parental wild-type (A) or serially ACE2-enriched cells (B) subsequently targeted by CRISPR-mediated single gene disruption of the indicated target gene or a nontargeting control. Candidate genes associated with positive or negative regulation of ACE2 in the CRISPR screen are highlighted in blue and red, respectively. ACE2-positive gates were defined on control unstained cells and the proportion of ACE2-positive cells for each population is displayed for each of 3 independent biologic replicates. Error bars depict standard deviation. Source data and statistical analysis are provided in Supplemental Table 7.

### ACE2 modifiers are enriched for regulators of gene expression, functional networks, and viral host factors

We next analyzed the 35 ACE2 modifiers identified in our CRISPR screens for enrichment in annotated gene ontologies and protein-protein interactions. We found the greatest enrichment for several molecular functions involved in gene expression, including transcriptional regulation, chromatin binding, and DNA binding (Figure 3A). ACE2 modifiers were significantly enriched (*p* <10^−12^) for annotated protein-protein interactions in the STRING database (Figure 3B) (Szklarczyk et al., 2019).

**Figure 3.**
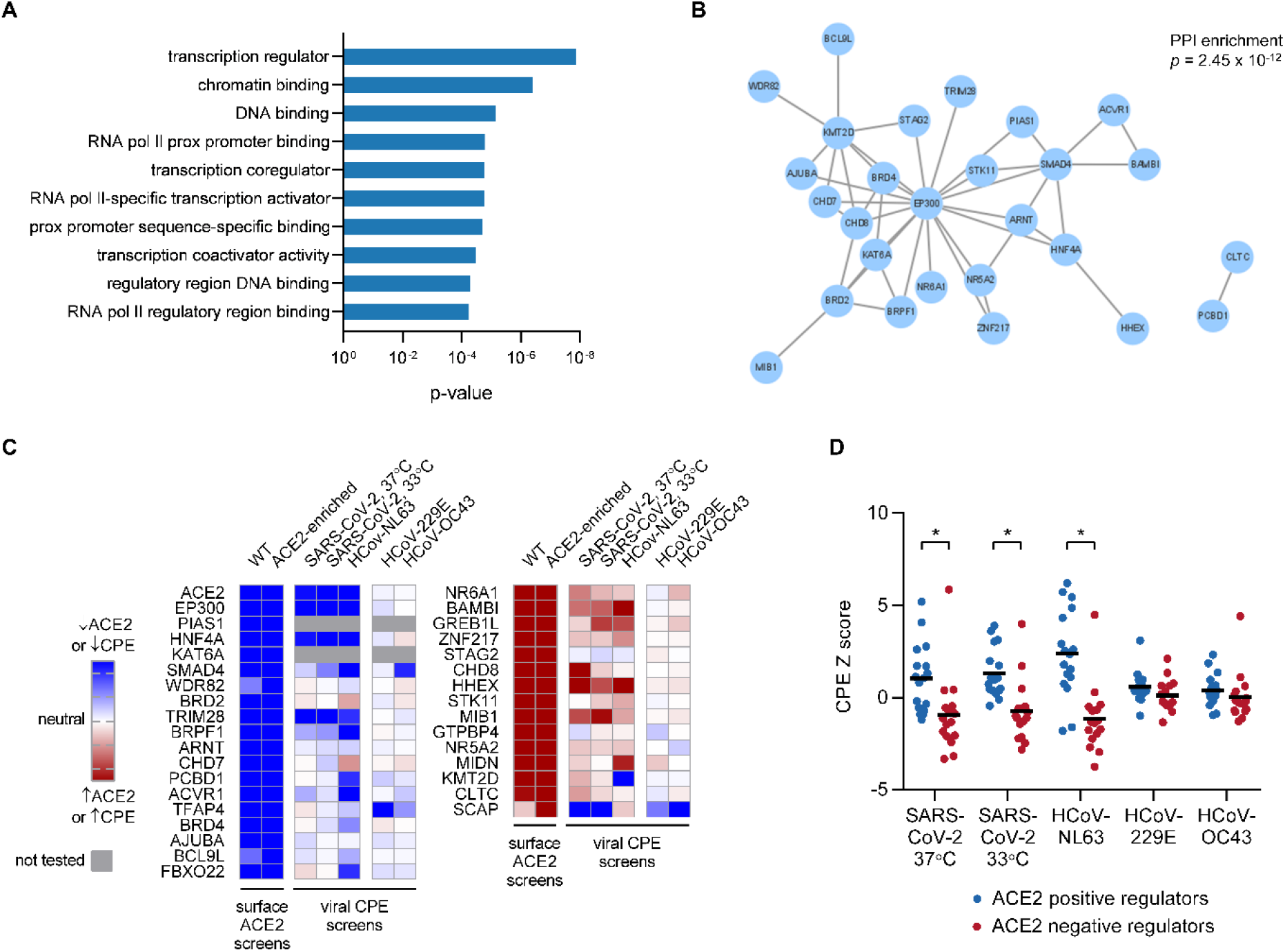
Analysis of ACE2 modifiers. (A) Top 10 molecular function ontologies enriched in genes identified as ACE2 modifiers relative to all genes tested in the focused CRISPR library. (B) Network analysis of ACE2 modifiers identified in the secondary CRISPR screen. Significance testing for the number of detected protein-protein interactions relative to a randomly selected gene set was calculated by STRING. (C) Heat map of log-transformed p-values for each ACE2 modifier identified in this study in comparison to scores from a reported study of cytopathic effect on HuH7.5 cells with the indicated coronaviruses (Schneider et al., 2021). (D) Comparison of CPE effect in Schneider et al for groups of genes identified in this study as ACE2 positive or negative regulators. Asterisks indicates p<0.001. A significant difference is observed for only those viruses whose cellular entry is ACE2-mediated.

We cross-referenced the data from our ACE2 CRISPR screen in HuH7 cells to that from a recently reported genome-wide CRISPR screen of HuH7.5 cellular sensitivity to infection by SARS-CoV-2 or by other coronaviruses (Schneider et al., 2021). As expected, *ACE2* itself was among the top genes identified both as a positive regulator of ACE2 surface abundance in HuH7 cells in our screen and as a proviral host factor for both SARS-CoV-2 and another ACE2-dependent coronavirus, HCoV-NL63, but not for other coronaviruses, HCoV-229E and HCoV-OC43, that use alternative cellular receptors (Figure 3C) (Hulswit et al., 2019; Yeager et al., 1992). Overall, among the genes we identified as ACE2 regulators, those that promoted ACE2 abundance in our screen were more likely to sensitize HuH7.5 cells to SARS-CoV-2 and HCoV-NL63 infection, while those that repressed ACE2 abundance were more likely to confer resistance to SARS-CoV-2 and HCoV-NL63 infection (Figure 3C, D). A similar correlation for ACE2 modifiers with cytopathic effect was not observed for HCoV-229E and HCoV-OC43 (Figure 3D). These findings support the relevance of our ACE2 screen to viral infection and suggest that even modest changes in ACE2 expression may influence cellular susceptibility to viral cytopathic effect.

### Cholesterol regulatory genes that influence coronavirus infection do not influence ACE2 surface abundance

Multiple CRISPR screens have implicated cholesterol regulation as an important mediator of host cell interactions with SARS-CoV-2 (Daniloski et al., 2021; Hoffmann et al., 2021; Schneider et al., 2021; Wang et al., 2021). In addition to canonical SREBP regulators, we also noted the identification of several genes in these studies that we had recently identified in a screen for regulators of low-density lipoprotein (LDL) uptake (Emmer et al., 2021). These included *RAB10* and multiple components of the exocyst complex (Hoffmann et al., 2021; Schneider et al., 2021; Wang et al., 2021). We therefore systematically examined the overlap among these CRISPR screens for LDL uptake, viral infection, and ACE2 abundance. We observed that gene disruptions which reduced LDL uptake were also more likely to confer cellular resistance to SARS-CoV-2 both at 37°C and at 33°C (Figure S3A). By contrast, disruption of these same genes was not associated with a significant change in ACE2 abundance in either of our screens of wild-type or ACE2-enriched HuH7 cells (Figure S3B). Further supporting an ACE2-independent effect on viral infection, positive regulators of LDL uptake did not influence cellular sensitivity to HCoV-NL63 (Figure S3A). Among these, disruption of the canonical SREBP regulators *SCAP, MBTPS1*, and *MBTPS2* conferred resistance to SARS-CoV-2 infection, with a similar effect for infection with the ACE2-independent HCoV-OC43 and HCoV-229E but not the ACE2-dependent HCoV2-NL63 (Figure S3C). These findings suggest that cholesterol regulatory host factors likely influence SARS-CoV-2 infection through an ACE2-independent mechanism.

### Regulation of ACE2 by *SMAD4, EP300, PIAS1*, and *BAMBI* is mediated at the mRNA level

Among the genes with the largest functional influence on ACE2 abundance in our screen and in our single gene validation experiments were *SMAD4* and multiple genes previously associated with SMAD4 signaling. These include *EP300*, encoding a histone acetyltransferase that is recruited by SMAD complexes to function as a coactivator for target genes (Feng et al., 1998; Janknecht et al., 1998); *PIAS1*, encoding an E3 sumo ligase whose substrates include SMAD4 and which has been shown to modulate SMAD4-dependent TGF-β signaling (Liang et al., 2004); and *BAMBI*, encoding a decoy receptor that negatively regulates TGF-β signaling through SMAD4 (Onichtchouk et al., 1999). To establish the level of regulation for ACE2 surface protein by these modifiers, we analyzed HuH7 cells individually targeted for each gene by CRISPR. To distinguish between changes in cell surface ACE2 being due either to a change in total cellular levels or to altered trafficking of ACE2, we quantified total cellular ACE2 protein abundance both by immunoblotting of cell lysates (Figure 4A) and by flow cytometry of permeabilized cells (Figure 4B). In all cases, changes in total cellular ACE2 abundance were consistent with changes in surface ACE2 abundance. Immunofluorescence microscopy of permeabilized cells likewise demonstrated alterations in total ACE2 abundance with no striking redistribution of ACE2 in CRISPR-targeted cells (Figure 4C). Quantitative RT-PCR demonstrated changes in *ACE2* mRNA levels in CRISPR-targeted cells that were consistent with the changes in surface-displayed ACE2 protein in these cells (Figure 4D). Together, these findings indicate that ACE2 regulation by *SMAD4, EP300, PIAS1*, and *BAMBI* in HuH7 cells is mediated at the mRNA level.

**Figure 4.**
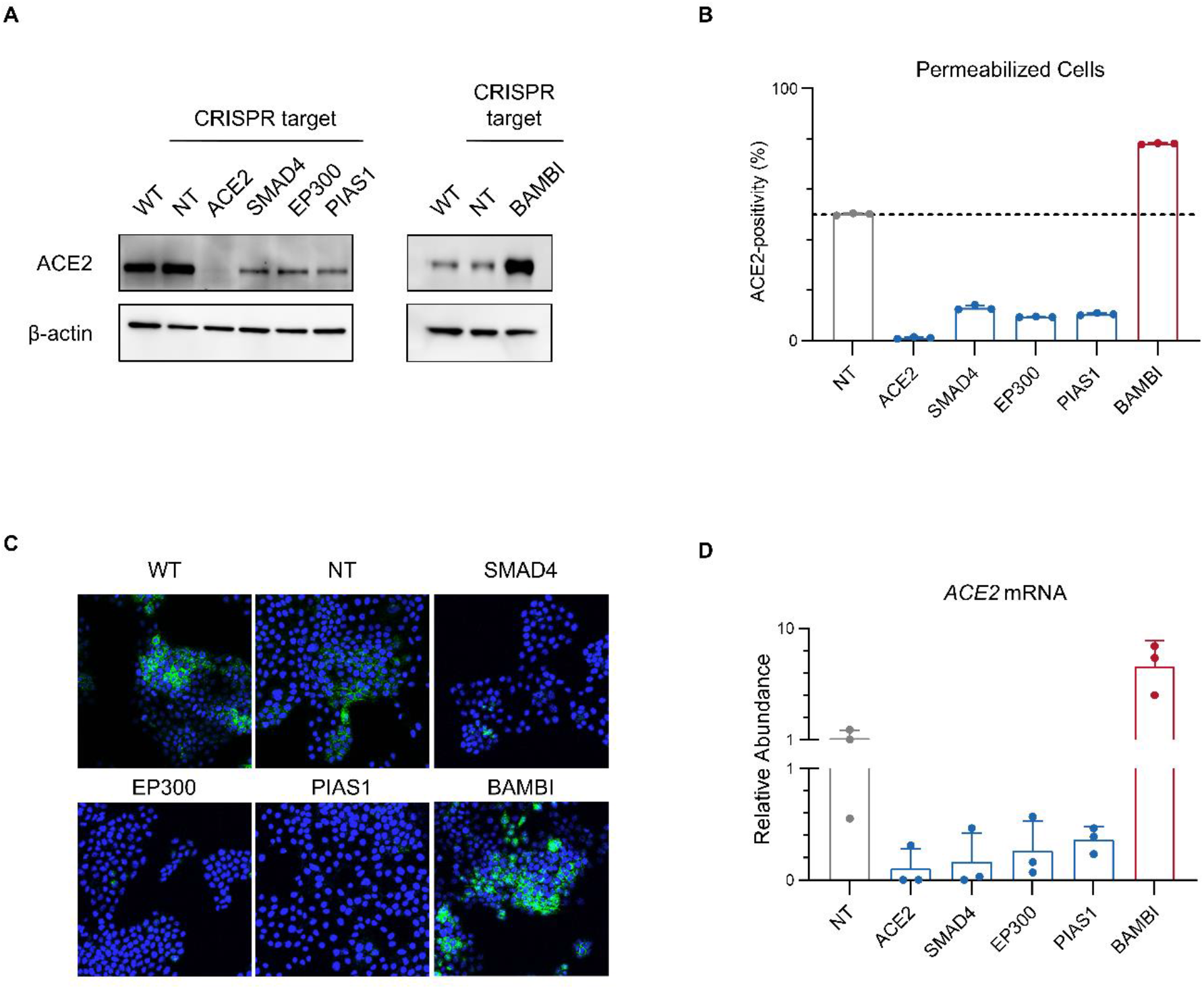
Regulation of ACE2 by SMAD4, EP300, PIAS1, and BAMBI is mediated at the mRNA level. (A) Immunoblotting of lysates collected from HuH7 cells either untreated or targeted by CRISPR with a gRNA against the indicated gene or a nontargeting (NT) control sequence. (B) Proportion of cells exhibiting ACE2 staining above background upon permeabilization with 0.1% saponin. (C) Immunofluorescence microscopy of ACE2 staining in HuH7 cells either untreated (WT) or targeted by CRISPR with a gRNA against the indicated gene or a nontargeting (NT) control sequence. (D) Quantification of relative ACE2 mRNA levels in the indicated cell lines by qRT-PCR of ACE2 mRNA, normalized to a panel of control transcripts.

### ACE2 modifiers alter cellular susceptibility to SARS-CoV-2 infection

We next analyzed HuH7 cells with CRISPR-mediated disruption of ACE2 modifiers for their sensitivity to SARS-CoV-2 infection. We found that those CRISPR-targeted cells with decreased ACE2 expression (*SMAD4, EP300*, and *PIAS1*) also exhibited a decrease in both the proportion of infected cells by SARS-CoV-2 nucleocapsid protein immunofluorescence (Figure 5A) and in viral infectivity by TCID50 (Figure 5B) at 48 hours post-infection. By contrast, cells with increased ACE2 expression (*BAMBI*) exhibited an increase in the proportion of SARS-CoV-2-infected cells and viral infectious titers. These findings, together with the correlation of results from our ACE2 screen and the HuH7.5 SARS-CoV-2 screen (Schneider et al., 2021), support the relevance of our identified ACE2 modifiers to SARS-CoV-2 infection.

**Figure 5.**
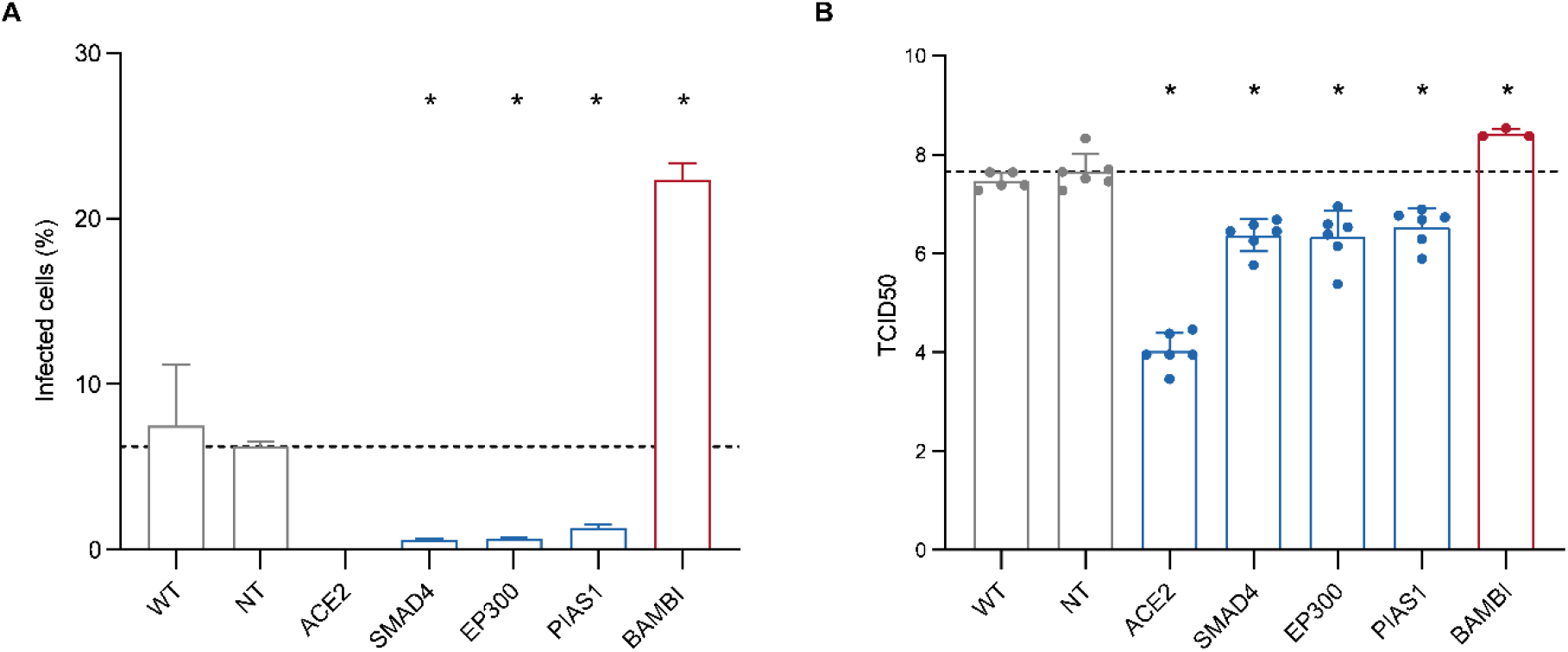
ACE2 regulators influence cellular sensitivity to SARS-CoV-2 infection. (A) Percentage of either wild-type or CRISPR-targeted cells with positive staining for SARS-CoV-2 nucleocapsid protein at 2 days post-infection with SARS-CoV-2 at MOI 1. An average of 214 fields of view (range 151 - 295) were analyzed for each condition over 2 independent biologic replicates. Error bars indicate standard deviation. (B) Infectious titers of cellular supernatants collected at 2 days post-infection with SARS-CoV-2 at MOI 1, plotted as log10(TCID50) with error bars indicating standard deviation. For both experiments, asterisks indicate *p* < 0.05 in comparison to nontargeting control cells, as calculated by one-way ANOVA with Dunnett correction for multiple comparison testing.

## DISCUSSION

CRISPR screening is a powerful forward genetic tool for the unbiased and high-throughput functional interrogation of the human genome. This approach has been successful in the identification of host factors involved in the molecular pathogenesis for many viruses (Puschnik et al., 2017), including SARS-CoV-2 (Baggen et al., 2021; Daniloski et al., 2021; Gordon et al., 2020; Heaton et al., 2020; Hoffmann et al., 2021; Schneider et al., 2021; Wang et al., 2021; Wei et al., 2021). For most identified host factors, however, the molecular basis for interaction with SARS-CoV-2 remains unclear. Consistently, these screens have lent further support to the central importance of ACE2 in SARS-CoV-2 infection, with guide RNAs targeting *ACE2* among those conferring the greatest resistance to cytopathic effect. Our current study, enabled by our recently reported identification of a sensitive and specific platform for ACE2 flow cytometry (Sherman and Emmer, 2021), complements the former studies to systematically probe the genetic regulators of ACE2 surface abundance in HuH7 cells.

An important caveat of our study is the focused nature of our library, limited to high-resolution functional testing of 833 candidate genes. Our initial attempts at genome-wide screening did nominate candidate ACE2 regulators that were included in our focused library, but these pilot studies were limited by an experimental bottleneck at the cell sorting stage with inadequate depth of coverage to support definitive conclusions about each gene tested. Nevertheless, for those genes interrogated by the focused library, several observations support a high degree of confidence in the functional significance of each gene to ACE2 surface abundance. First, the internal control *ACE2*-targeting gRNAs were clearly depleted in ACE2-positive cells. Second, the genes that were identified as regulating ACE2 abundance exhibited a robust statistical enrichment or depletion. Third, we observed a very high degree of concordance in the degree of enrichment or depletion observed for each gene between the independent screens of wild-type or ACE2-enriched HuH7 cells. Fourth, identified genes were highly enriched for related functional annotations and protein-protein interactions and exhibited significant overlap with previously identified modulators of HuH7.5 cellular susceptibility to ACE2-dependent coronaviruses. Finally, of the screen hits we selected for single gene validation testing, the vast majority (18 of 21) were confirmed to regulate ACE2 surface abundance.

Given the sensitivity of our screens to detect subtle influences on ACE2 abundance, a number of negative findings are noteworthy. Cholesterol regulatory genes in the SREBP pathway have been implicated in SARS-CoV-2 infection, but we did not detect an association with ACE2 abundance either by focused analysis of canonical SREBP regulators or by systematic comparison to all HuH7 LDL uptake regulators identified in our previously reported genome-wide CRISPR screen (Emmer et al., 2021). These findings are consistent with the association of SREBP regulators with ACE2-independent coronaviruses and the lack of association with ACE2-dependent HCoV-NL63. Angiotensin receptor blockade has been postulated to increase surface ACE2 expression, raising concern for potential increased COVID-19 risk for patients on ACE inhibitors or angiotensin receptor blockers. We did not however detect any association of *AGTR1* or *AGTR2* with ACE2 abundance. We also did not observe an effect for disruption of the Ang[1-7] receptor *MAS1*, arguing against feedback regulation of this axis in these cells. Although estrogen regulation of ACE2 expression has been proposed to mediate the sexual dimorphism of COVID-19 susceptibility, and estrogen receptor binding sites have been identified near the ACE2 locus (Barker and Parkkila, 2020), we did not detect an effect of *ESR1* or *ESR2* disruption on ACE2 abundance. We also did not detect a large effect for any candidate genes in proximity to variants associated with COVID-19 susceptibility by GWAS. Each of these negative results should be interpreted with caution, however, both because of the potential cell type specificity of ACE2 regulation and the intrinsic limitation of CRISPR screens for functionally redundant genes, essential genes, or compensatory mechanisms in mutant cells.

SMAD4, which we identified as a positive regulator of ACE2 gene expression, is a common mediator of TGF-β signaling (Zhao et al., 2018). In the canonical TGF-β signaling pathway, ligand binding signals through receptor-regulated SMADs that complex with SMAD4 to trigger its translocation into the nucleus and binding to specific DNA regulatory elements. Identification of the upstream signals driving SMAD4-dependent ACE2 expression in HuH7 cells is complicated by the diversity of over 40 TGF-beta superfamily receptors and a variety of ligands including bone morphogenic proteins, nodal, and activin. Our screen did identify *ACVR1*, encoding activin A receptor type 1, as a positive regulator of ACE2, and this gene was similarly identified as proviral for SARS-CoV-2 and HCoV-NL63 in HuH7.5 cells (Schneider et al., 2021). The limited scope of our screen, however, precludes a comprehensive cataloguing of other potential upstream mediators of SMAD4-dependent ACE2 expression. Consistent with our results, a recent study identified ChIP-seq peaks for SMAD4 in intestinal epithelial cells that overlapped putative enhancer regions near the murine *Ace2* locus, and furthermore detected reduced *Ace2* intestinal transcript levels upon tissue-specific *Smad4* deletion (Chen et al., 2021).

Our findings are consistent with a growing body of evidence supporting the cell type specificity of genetic interactions relevant to SARS-CoV-2 infection. Although we observed significant overlap in the ACE2 modifiers identified in our screens with recently identified SARS-CoV-2 infection modifiers in HuH7.5 cells (Schneider et al., 2021), minimal overlap was observed with a similar screen of Vero E6 cells (Wei et al., 2021). In the latter study, single gene disruption of *SMAD4* in Calu-3 cells similarly was not associated with SARS-CoV-2 resistance. As expected, we observed little overlap in our identified ACE2 modifiers with other genome-wide SARS-CoV-2 screens that used cell lines engineered with ectopic expression of ACE2 (Baggen et al., 2021; Daniloski et al., 2021; Wang et al., 2021). For those SARS-CoV-2 screens using cells with endogenous ACE2 expression, it is possible that this discordance may be attributable to technical differences between the screens. However, a cell type-specific network regulating SARS-CoV-2 entry is also supported by small molecule inhibitor studies (Dittmar et al., 2021) and by the tissue-specific patterns of ACE2 promoter usage (Pedersen et al., 2013).

In summary, we have applied a functional genomic approach to dissect the regulatory networks of ACE2 protein expression in HuH7 cells. We have identified many previously unrecognized genetic modifiers of ACE2 expression and a putative mechanism for genes previously implicated in SARS-CoV-2 infection. For *SMAD4, EP300, PIAS1*, and *BAMBI*, we established their level of regulation on ACE2 and their influence on SARS-CoV-2 infection. These findings clarify the molecular determinants of ACE2 expression and nominate pathways for host-targeted therapeutic development.

## Acknowledgments

We thank all individuals involved in this multidisciplinary collaboration conducted under the challenging circumstances of the COVID-19 global pandemic. This research was supported by the National Institutes of Health K08-HL148552 (BTE), the University of Michigan Frankel Cardiovascular Center (BTE), the Michigan Institute for Clinical and Health Research training grant LT1 (CM), the Marie-Slodowska Curie global fellowship GA – 841247 (CM), and the University of Michigan Biological Scholars Program (CEW).

## Author Contributions

EJS and BTE conceived the project. EJS, SEG, CJW, AWT, JZS, CEW, and BTE designed the CRISPR library. EJS, CM, and BTE performed the CRISPR library synthesis and screening. EJS, CM, VTT, TGK, and BTE performed screen follow-up including bioinformatics analysis, arrayed validation, and analysis of single gene CRISPR-targeted cell lines. CM performed all BSL3 work. All authors contributed to data analysis and manuscript review. EJS and BTE wrote the manuscript with input from all authors.

## MATERIALS AND METHODS

### Genome-wide ACE2 CRISPR screen

Wild-type or serially ACE2-enriched HuH7 cells were independently screened. For each of 4 biologic replicates for each cell line, a total of ∼100 million cells were transduced with the GeCKOv2 library (Sanjana et al., 2014) at an MOI of ∼0.3. Puromycin was added at a concentration of 3 µg/mL at day 1 post-transduction and maintained until selection of control uninfected cells was complete. Cells were passaged as needed to maintain logarithmic phase growth with total cell number maintained above a minimum of 25 million cells at all stages of the screen. At 14 days post-transduction, a total of ∼200 million cells were harvested and stained for surface ACE2 abundance as previously described with ACE2 antibody (R&D Systems #MAB9332) at 1:50 dilution in FACS buffer (PBS supplemented with 2% FBS) and Alexa Fluor 647-conjugated goat anti-rabbit IgG secondary antibody (AlexaFluor647 goat anti-rabbit IgG (Fisher #A32733) at 1:500 dilution in FACS buffer. Flow cytometry gates were defined for cell viability by exclusion of SYTOX Blue Dead Cell Stain (Fisher, S34857) and for ACE2 expression by comparison to unstained or ACE2-targeted control cells. For the screen of wild-type HuH7 cells, the ∼3-5% of the total cells that were positive for ACE2 staining (as defined by gating of unstained cells) were collected. For the screen of ACE2-enriched HuH7 cells, the brightest 10% ACE2-positive cells were collected. For both screens, a gate of the median 10% of ACE2-negative cells were also collected. Genomic DNA was extracted and gRNA sequences amplified and sequenced as previously described (Emmer et al., 2021).

### CRISPR screen analysis

FASTQ files were processed by PoolQ (Broad Institute; https://portals.broadinstitute.org/gpp/public/software/poolq) to map individual sequencing reads to reference gRNA sequences with deconvolution by barcode. Cumulative distribution functions of gRNA representation were generated by plotting normalized read counts of each gRNA against its relative rank for a given barcode. Individual gRNA-level and aggregate gene-level enrichment analysis was performed using MAGeCK(Li et al., 2014). Q-Q plots were generated by plotting log-transformed observed p-values (calculated by MAGeCK gene-level analysis) against expected p-values (determined by the relative rank of each gene among the library). Genes were considered screen hits if they were identified with a MAGeCK-calculated false discovery rate < 0.05 and absolute log2 fold-change > 1 in either analysis of HuH7 wild-type or ACE2-enriched cells. Enrichment of molecular functions among secondary screen hits relative to all genes in the secondary library was performed using GOrilla(Eden et al., 2009). Gene network analysis of secondary screen hits was performed using the STRING database (Szklarczyk et al., 2019) with default settings and visualized with Cytoscape (Shannon et al., 2003). Heatmaps were generated with log-transformed p-values from our study and from Schneider et al.(Schneider et al., 2021) using GraphPad Prism v9.1.0.

### Design and synthesis of secondary CRISPR library

Candidate genes were selected from (i) top candidate genes identified in our primary genome-wide CRISPR screen for ACE2 surface abundance; (ii) genes identified in CRISPR screens as candidate modifiers of SARS-CoV-2 cytopathic effect (Daniloski et al., 2021; Gordon et al., 2020; Heaton et al., 2020; Hoffmann et al., 2021; Wang et al., 2021; Wei et al., 2021); (iii) genes whose expression was correlated with ACE2 surface abundance in HuH7 cells (Sherman and Emmer, 2021); (iv) genes in proximity to loci associated with COVID-19 susceptibility by GWAS (described below); (v) candidate genes identified in our own pilot CRISPR screens of ACE2 surface abundance in ACE2-overexpressing HEK293T cells or Caco2 cells or SARS-CoV-2 cytopathic effect in Caco2 cells; and (vi) hypothesis-driven manually selected genes. For each gene, a total of 15 optimized gRNA sequences were identified using the Broad Genetic Perturbation Platform (Sanson et al., 2018). For identification of COVID-19 GWAS loci, data freeze 4 of the COVID-19 Host Genetics Initiative was used (Ganna, 2021). Lead SNPs were selected from the analysis of hospitalized COVID-19 patients relative to population controls. For each indicated SNP, genes were selected based on their physical proximity to the SNP and by their Polygenic Priority Score (Weeks et al., 2020) (Supplemental Table 2). Flanking sequences were appended to facilitate PCR amplification and oligonucleotides were synthesized by CustomArray (Bothell, WA). DNA assembly of the secondary library plasmid pool was performed with 125 ng of PCR amplicon and 825 ng of BsmBI-digested pLentiCRISPRv2 in a total reaction volume of 100 µL with HiFI DNA Assembly Mix (NEB) for 30 min at 50°C. Assembly products were purified with a QIAquick PCR purification kit (Qiagen, Hilden, Germany) and 5 electroporations were performed into Endura electrocompetent cells (Lucigen, Middleton WI) and plated onto 24.5 cm^2^ LB-agar plates. After 14 hr at 37°C, bacteria were harvested and plasmid DNA purified with an EndoFree Plasmid Maxi kit (Qiagen). Dilution plates of electroporated cells confirmed a colony count of >100X relative to the size of the gRNA library and representation was confirmed by sequencing on an Illumina MiSeq with gRNA mapping and cumulative distribution function analysis as described above. Lentiviral stocks were generated by cotransfection of the lentiviral plasmid pool with psPAX2 and pVSVG into HEK293T cells, harvesting of supernatants, and titering of virus stocks as previously described (Emmer et al., 2018).

### Arrayed validation of ACE2 modifiers

For each gene tested, a single gRNA was selected from the 15 in the library based on its degree of enrichment or depletion in the secondary screens. Each individual gRNA was ligated into *BsmBI*-digested pLentiCRISPRv2(Sanjana et al., 2014) and lentiviral stocks generated and titered as previously described(Emmer et al., 2018). HuH7 wild-type and ACE2-enriched cells were transduced in parallel with each lentiviral construct, treated with puromycin 3 µg/mL until no surviving cells remained among control non-transduced cells, and passaged to remain logarithmic phase growth. Surface ACE2 abundance was quantified at day 14 post-transduction by flow cytometry as previously described (Sherman and Emmer, 2021).

### Analysis of ACE2 mRNA and protein

Immunoblotting of HuH7 RIPA lysates was performed as previously described (Sherman and Emmer, 2021) with antibodies against ACE2 (GeneTex, Irving CA, #GTX01160, 1:1000), β-actin (Santa Cruz Biotechnology, Dallas TX, #sc-47778, 1:5000), SMAD4, EP300, PIAS1, and BAMBI. Permeabilization prior to ACE2 flow cytometry was performed by incubating cells prior to blocking in 0.1% saponin in FACS buffer for 10 min, with subsequent ACE2 staining performed as described above. Immunoflourescence microscopy of ACE2 was performed on cells seeded in 35 mm poly-D lysine-coated glass bottom dishes (MatTek, Ashland MA, P35GC-1.5-14-C) as previously described (Liang et al., 2004). Quantification of ACE2 transcript levels was performed by preparing total RNA from 2-4×10^6^ cells for each sample using the RNeasy Plus Micro kit (Qiagen, Hilden Germany, #74034). For qRT-pCR, cDNA was prepared using the SuperScript III first-strand synthesis kit (Thermo Fisher, #18080051), amplified with indicated primers using the Power SYBR Green PCR Master Mix (Thermo Fisher, # 4367659), and analyzed by QuantStudio 5 Real-Time PCR (Thermo Fisher).

### SARS-CoV-2 infection assays

SARS-CoV-2 WA1 strain was obtained by BEI resources and was propagated in Vero E6 cells. Lack of genetic drift of our viral stock was confirmed by deep sequencing. Viral titers were determined by TCID50 assays in Vero E6 cells (Reed and Muench method) by microscopic scoring. All experiments using SARS-CoV-2 were performed at the University of Michigan under Biosafety Level 3 (BSL3) protocols in compliance with containment procedures in laboratories approved for use by the University of Michigan Institutional Biosafety Committee (IBC) and Environment, Health and Safety (EHS). For the immunofluorescence-mediated assay, 384-well plates (Perkin Elmer, 6057300) were seeded with HuH7 cells at 3000 cells per well and allowed to adhere overnight. Plates were then transferred to BSL3 containment and infected with SARS-CoV-2 WA1 at a multiplicity of infection (MOI) of 1. Two days post-infection, cells were fixed with 4% PFA for 30 minutes at room temperature, permeabilized with 0.3% Triton X-100, and blocked with antibody buffer (1.5% BSA, 1% goat serum and 0.0025% Tween 20). The plates were then sealed, surface decontaminated, and transferred to BSL2 for staining with antibody against SARS-CoV-2 nucleocapsid protein (Antibodies Online, Cat# ABIN6952432) overnight at 4 ?C followed by staining with Alexa Fluor 647-conjugated secondary antibody (goat anti-mouse, Thermo Fisher, A21235) and DAPI (Thermo FIsher). Plates were imaged with Thermo Fisher CX5 high content microscopes with a 10X/0.45NA LUCPlan FLN objective and analyzed with a Cell Profiler pipeline. Percentage of infected cells was calculated as previously described as N-protein positive cells versus DAPI-positive (Mirabelli et al., 2020). For the infectivity assay, HuH7 were seeded at 3 × 10^5^ cells per well in a 12-well plate and allowed to adhere overnight. The next day, cells were infected with SARS-CoV-2 WA1 at a MOI of 1 at 37°C for 1hr. Viral inoculum was removed by serial washing (three times). Two days post-infection cells were harvested by scraping the monolayers and lysates were centrifuged at high speed for 5 minutes to allow the release of intracellular viral progeny in infected cells. Infectious titer of the supernatants was determined by TCID50 assay.

---

**Figure S1.**
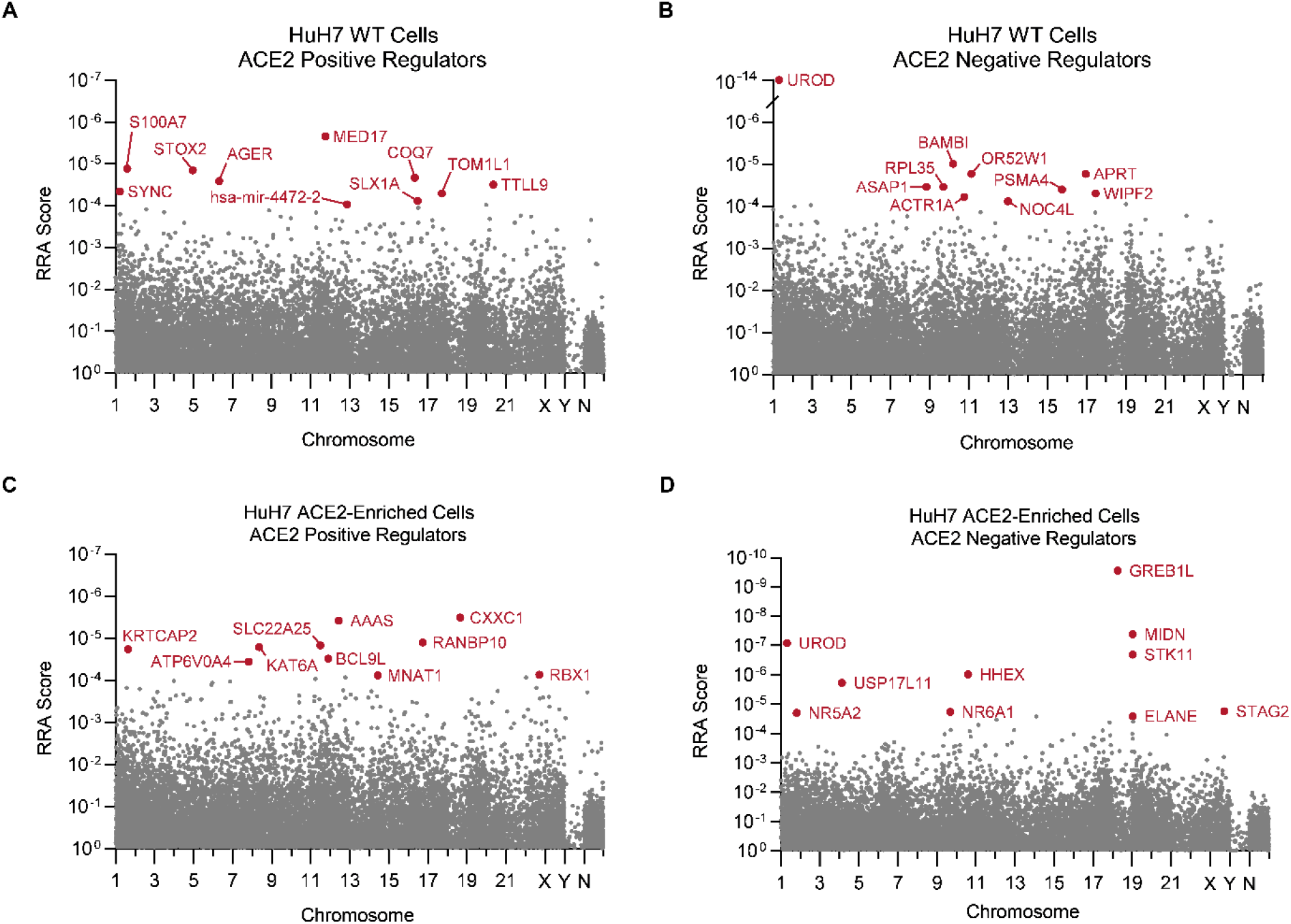
Primary genome-wide CRISPR screen for ACE2 modifiers. Manhattan plots demonstrating gene-level MAGeCK Robust Rank Aggregation (RRA) scores for each gene in the GeCKOv2 library plotted according to its chromosomal transcription start site. N = nontargeting controls. Plots are displayed for both positive regulation (A, C; gRNAs depleted in ACE2-positive relative to ACE2-negative populations) and negative regulation (B, D; gRNAs enriched in ACE2-positive relative to ACE2-negative populations) for independent screens of HuH7 wild-type cells (A, C) and HuH7 ACE2-enriched cells (B, D). The top 10 genes for each analysis are highlighted in red.

**Figure S2.**
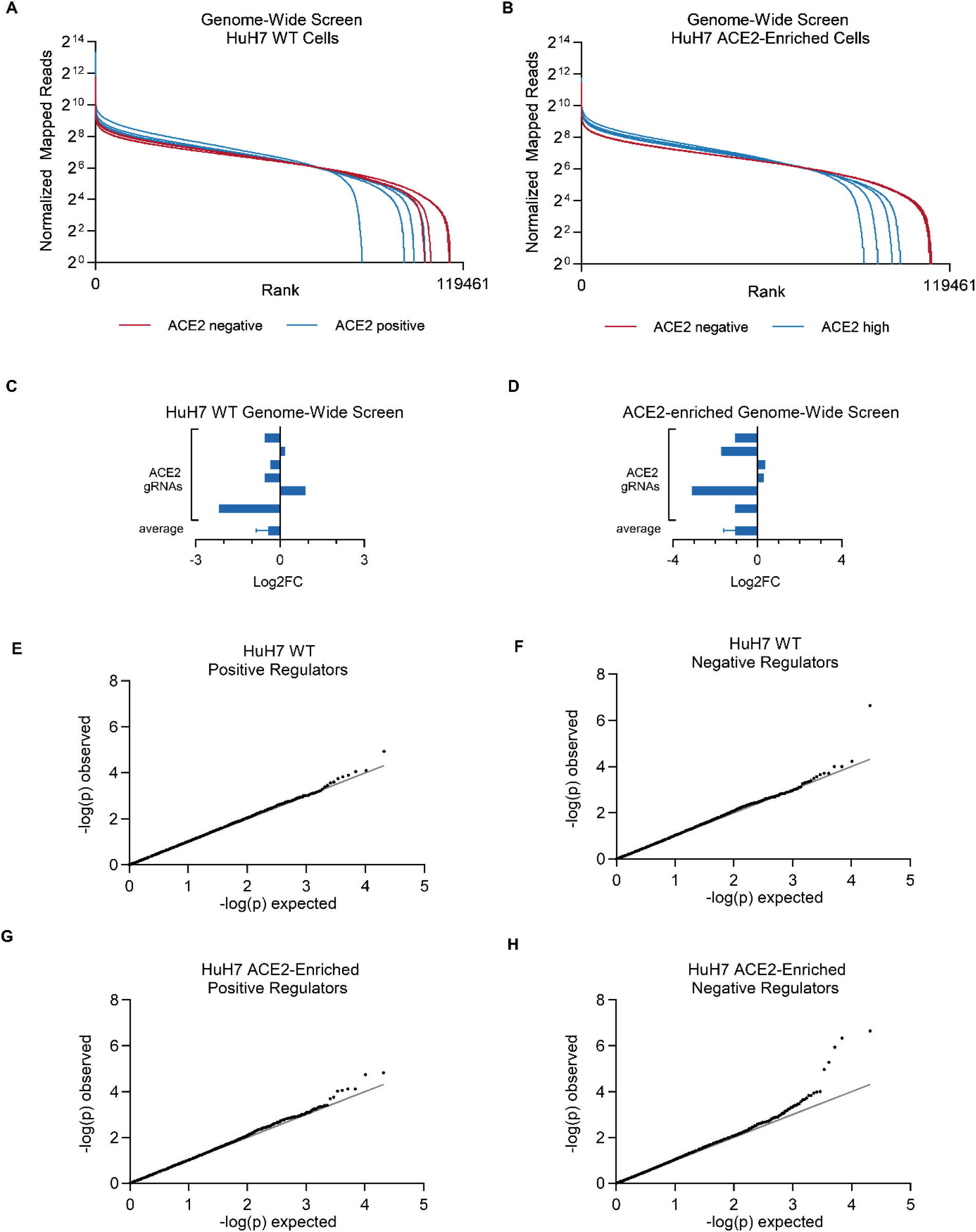
Quality analysis of genome-wide ACE2 CRISPR screens. (A-B) Cumulative distribution functions of normalized read counts for each gRNA in ACE2-sorted populations in 4 independent biologic replicates of the genome-wide primary CRISPR screen of wild-type (A) and ACE2-enriched (B) HuH7 cells. (C) Mean log2 fold-change of each individual gRNA targeting ACE2 in the genome-wide ACE2 CRISPR screens of wild-type (C) and ACE2-enriched (D) HuH7 cells. (E-H) Q-Q plots of observed versus expected –log(p) for positive (E,G) or negative (F,H) regulation of ACE2 surface abundance for every gene in the primary genome-wide CRISPR screen of HuH7 wild-type (E-F) or ACE2-enriched (G-H) cells. Observed p-values calculated by MAGeCK gene-level analysis.

**Figure S3.**
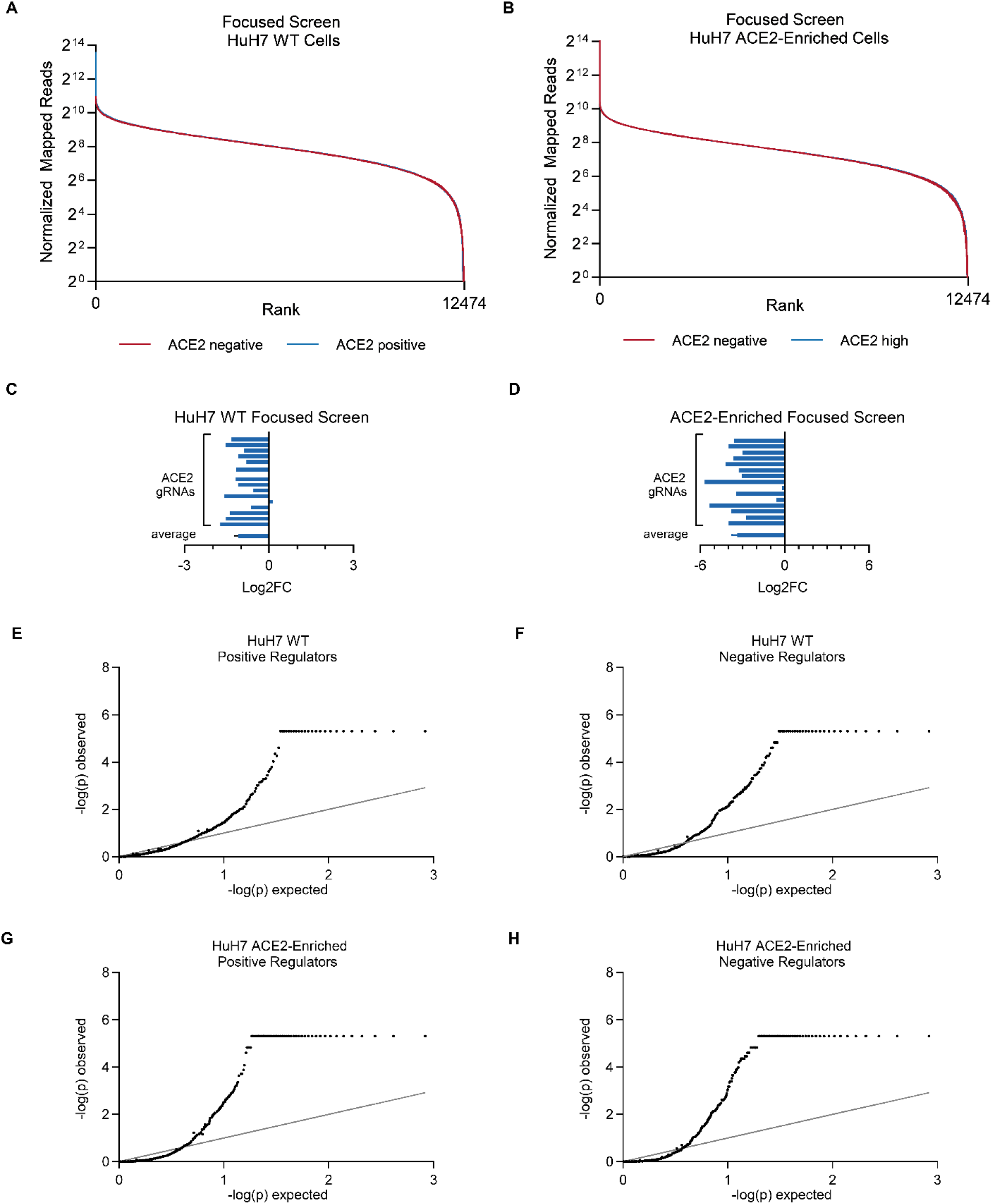
Quality analysis of focused ACE2 CRISPR screens. (A-B) Cumulative distribution functions of normalized read counts for each gRNA in ACE2-sorted populations in 3 independent biologic replicates of the focused secondary CRISPR screen of wild-type (A) and ACE2-enriched (B) HuH7 cells. (C-D) Mean log2 fold-change of each individual gRNA targeting ACE2 in the focused ACE2 CRISPR screens of wild-type (C) and ACE2-enriched (D) HuH7 cells. (E-H) Q-Q plots of observed versus expected –log(p) for positive (E,G) or negative (F,H) regulation of ACE2 surface abundance for every gene in the primary genome-wide CRISPR screen of HuH7 wild-type (E-F) or ACE2-enriched (G-H) cells. Observed p-values calculated by MAGeCK gene-level analysis.

**Figure S4.**
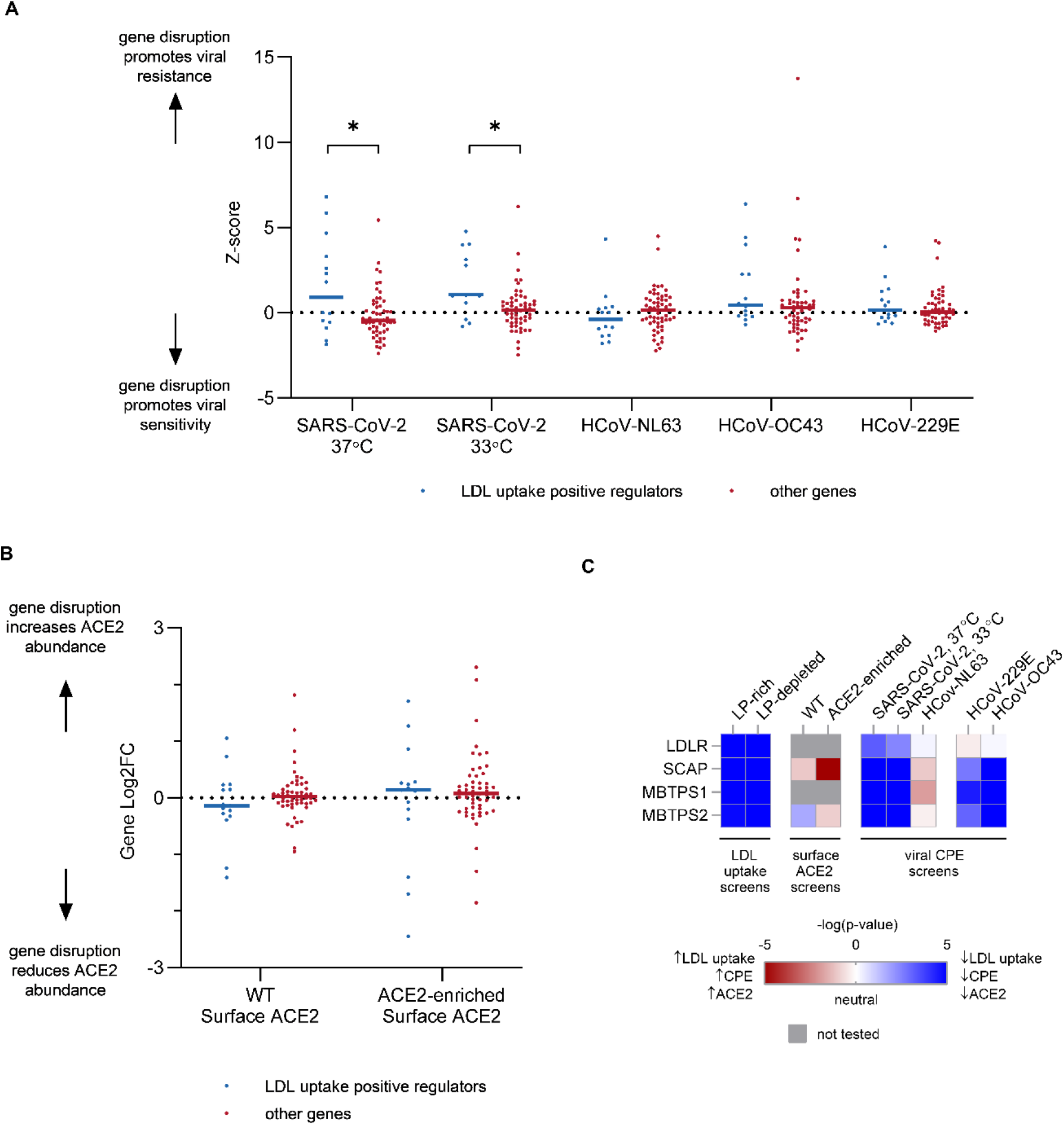
Comparison of CRISPR screen results for modifiers of HuH7 endocytosis, HuH7.5 SARS-CoV-2 infection, and HuH7 ACE2 surface abundance. (A) The subset of genes tested by high-resolution CRISPR screening for both LDL endocytosis and ACE2 surface abundance were divided into those whose disruption reduced LDL uptake (n=15) and those that did not (n=55). The Z-score for each gene within each group was then compared for each of the viral CPE screens reported by Schneider et al. Asterisks indicate p < 0.05 by Student’s t-test.(B) Genes were grouped as in (A) and compared for their association with surface ACE2 abundance in secondary screens of either HuH7 wild-type or ACE2-enriched cells. (C) Heat maps for screen results of canonical SREBP regulators LDLR, SCAP, MBTPS1, and MBTPS2.

**Figure S5.**
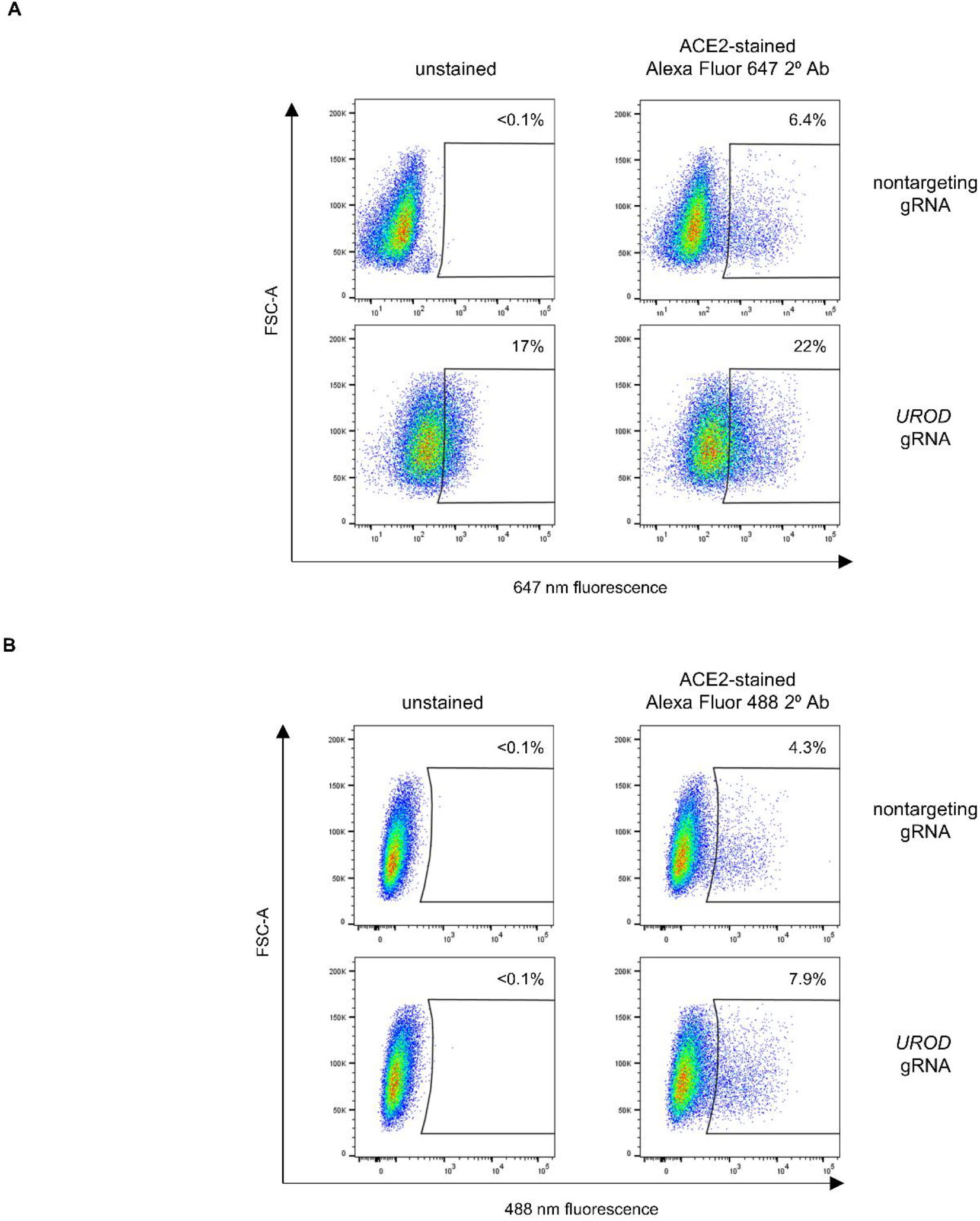
Disruption of *UROD* increases cellular autofluorescence. (A-B) FACS plots of fluorescence intensities of control and *UROD*-targeted HuH7 cells either unstained or incubated with an ACE2 antibody and a corresponding Alexa Fluor 488 (A) or Alexa Fluor 647 (B) conjugated secondary antibody.

**Figure S6.**
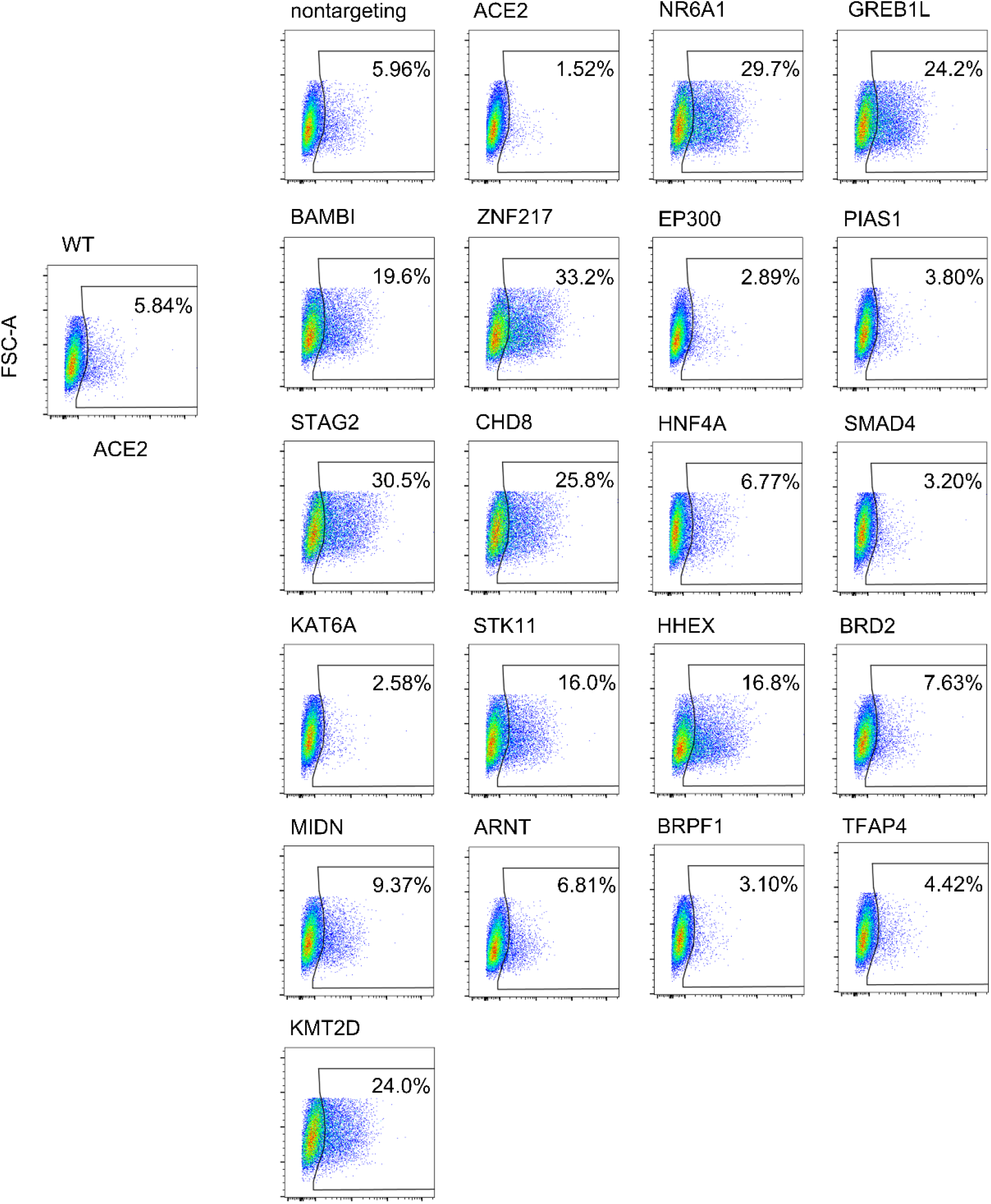
Arrayed validation of ACE2 modifiers on a HuH7 wild-type background. FACS plots of ACE2 staining relative to forward scatter area for untreated and single gRNA CRISPR-targeted HuH7 cells. Source data for a representative replicate of Figure S3 are displayed.

**Figure S7.**
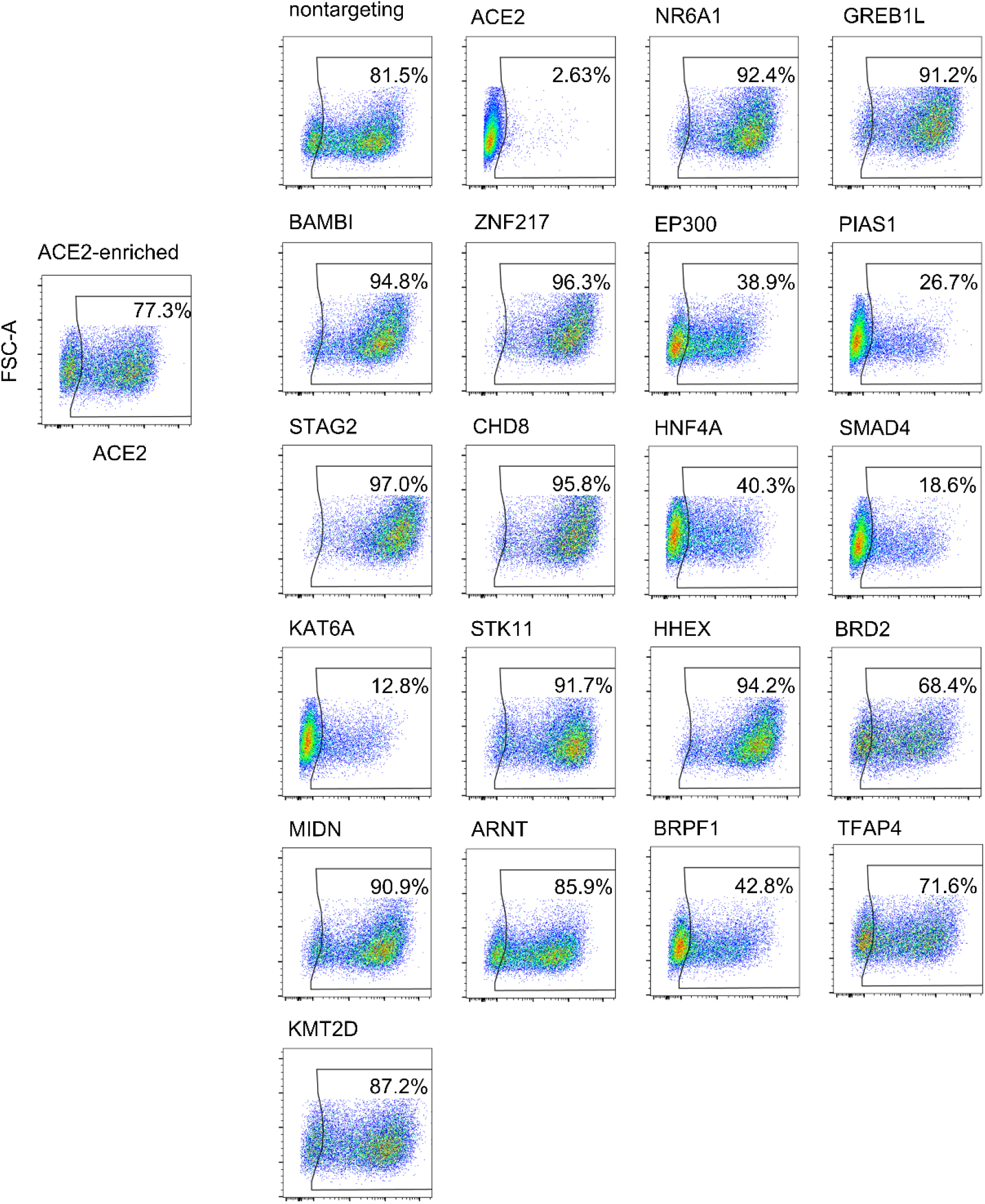
Arrayed validation of ACE2 modifiers on a HuH7 ACE2-enriched background. FACS plots of ACE2 staining relative to forward scatter area for untreated and single gRNA CRISPR-targeted HuH7 ACE2-enriched cells. Source data for a representative replicate of Figure S3 are displayed.

**Supplemental Table 1. CRISPR library design**. Primary sources and selection criteria are indicated for the gene lists that comprise the focused CRISPR library of potential modifiers.

**Supplemental Table 2. GWAS candidate gene identification**. Top-scoring SNPs associated with COVID-19 infection are listed along with candidate causal genes selected by either their physical proximity or Polygenic Prioritization Score.

**Supplemental Table 3. Genome-wide ACE2 CRISPR screen of HuH7 wild-type cells**. MAGeCK output for gene-level and individual gRNA-level enrichment or depletion in ACE2-positive cells relative to ACE2-negative cells. Negative log2-fold change and RRA scores indicate gRNA depletion in ACE2-positive cells (gene disruption associated with reduced ACE2 abundance).

**Supplemental Table 4. Genome-wide ACE2 CRISPR screen of HuH7 ACE2-enriched cells**. MAGeCK output for gene-level and individual gRNA-level enrichment or depletion in ACE2-high cells relative to ACE2-negative cells. Negative log2-fold change and RRA scores indicate gRNA depletion in ACE2-high cells (gene disruption associated with reduced ACE2 abundance).

**Supplemental Table 5. Focused CRISPR screen of HuH7 wild-type cells**. MAGeCK output for gene-level and individual gRNA-level enrichment or depletion in ACE2-positive cells relative to ACE2-negative cells. Negative log2-fold change and RRA scores indicate gRNA depletion in ACE2-positive cells (gene disruption associated with reduced ACE2 abundance).

**Supplemental Table 6. Focused ACE2 CRISPR screen of HuH7 ACE2-enriched cells**. MAGeCK output for gene-level and individual gRNA-level enrichment or depletion in ACE2-high cells relative to ACE2-negative cells. Negative log2-fold change and RRA scores indicate gRNA depletion in ACE2-high cells (gene disruption associated with reduced ACE2 abundance).

**Supplemental Table 7. Arrayed validation of ACE2 modifiers**. Individual replicate values of ACE2-positivity in populations either untreated (WT) or transduced with a single gene-targeted lentiCRISPR for the indicated gene or non-targeting (NT) control gRNA. Statistical testing was performed by Student’s t-test of 2-tailed distributions assuming equal variance.

